# FMRP activates the translation of large autism/intellectual disability proteins and stimulates N-end rule E3 ligase Poe/UBR4 production within puromycin-sensitive RNP particles

**DOI:** 10.1101/2020.06.27.174136

**Authors:** Ethan J. Greenblatt, Allan C. Spradling

**Affiliations:** Department of Biochemistry and Molecular Biology, University of British Columbia, 2350 Health Sciences Mall, Vancouver, British Columbia, V6T 1Z3 Canada; Howard Hughes Medical Institute Research Laboratories, Department of Embryology, Carnegie Institution for Science, 3520 San Martin Dr., Baltimore, Maryland 21218 USA

**Keywords:** Fmr1, FMRP, autism, RNP, translation, Poe/Ubr4

## Abstract

Mutations in Fmr1, encoding the RNA binding protein FMRP, are leading causes of intellectual disability, autism, and female infertility, but FMRP’s mechanism of action is controversial. In contrast to its previously postulated function as a translation repressor acting by stalling elongation, we recently found that FMRP activates translation initiation of large proteins in Drosophila oocytes up to ∼2-fold. We report here that FMRP’s function as a translational activator is conserved in the mammalian brain. Reanalysis of mouse cortex ribosome profiling data shows that translation of large proteins in Fmr1 mutants is down-regulated 2.0-1.2-fold; ribosome stalling appears not to influence FMRP target protein translation in either cortex or oocyte tissue. Consistent with an activator function, most FMRP targets are associated with clinical syndromes when reduced, but not when over-expressed. Fmr1-dependent translation of one target, the N-end rule E3 ligase Poe/UBR4, occurs in microscopically visible ribonucleoprotein particles. These “Poe particles” require FMRP for their formation, are distinct from P bodies, and depend on actively elongating ribosomes, as indicated by their dissolution following a brief puromycin treatment. N-end rule-mediated proteolysis via Poe/UBR4 restrains cell growth and limits MAPK signaling in nervous tissue. Thus, loss of FMRP reduces production of an important growth repressor.

## INTRODUCTION

Mutations in the Fmr1 gene, encoding the RNA binding protein FMRP, lead to fragile X syndrome (FXS) and fragile X primary ovarian insufficiency (FXPOI), causes of intellectual disability (ID), autism spectrum disorder (ASD), and premature ovarian failure (Hagerman et al. 2017; Sullivan et al. 2011). Neurons and oocytes control the translation and transport of mRNAs using ribonucleoprotein particles (RNPs), a subset of which contain FMRP (Barbee et al. 2006; Rosario et al. 2016; Antar et al. 2005). FMRP targets are non-randomly enriched in genes associated with ID and ASD disorders (Darnell et al. 2011; Ascano et al. 2012). FMRP binds to and potentially regulates the translation of hundreds of individual mRNAs (Darnell et al. 2011; Ascano et al. 2012). These targets encode proteins involved in a wide variety of biological processes, and include chromatin remodeling enzymes, microtubule adaptors, synaptic scaffolding proteins, and ubiquitin ligases (Darnell et al. 2011; Sawicka et al. 2019). How FMRP interacts with its target mRNAs, including ASD risk genes, and how FMRP-containing RNPs control target translation remain incompletely understood.

FMRP-deficient neurons exhibit an exaggerated form of activity-dependent neuronal plasticity known as mGluR5-dependent long-term depression (LTD) (Bear et al. 2004; Bear 2005), which has garnered considerable attention as a potential therapeutic target (Pop et al. 2014). FMRP has been proposed to attenuate LTD by inhibiting activity-dependent protein translation, as FMRP promotes ribosomal stalling during the elongation phase of translation *in vitro* (Darnell et al. 2011; Chen et al. 2014; Shah et al. 2020). Inconsistent with this model, however, is the finding by multiple groups that blocking activity-dependent translation with small molecule inhibitors does not rescue exaggerated mGluR5-dependent LTD in FMRP-deficient neurons (Hou et al. 2006; Park et al. 2008; Ronesi et al. 2012). Efforts to target the mGluR5 system as a therapeutic approach for fragile X syndrome have proven unsuccessful (Erickson et al. 2017).

Measurements of Drosophila oocyte protein production with ribosome profiling revealed that FMRP preferentially affects large proteins, and that it promotes rather than represses translation (Greenblatt and Spradling 2018). These divergent effects of FMRP in mice and Drosophila could be due to species and/or tissue differences. However, FMRP is highly conserved in structure and function between Drosophila and mouse, and earlier studies relied on *in vitro* systems to reconstitute FMRP’s proposed repressive activity. Moreover, long protein coding capacity is emerging as a common property shared by most mRNA targets regulated by Fmr1 in multiple systems (Greenblatt and Spradling 2018; Sears et al. 2019; Das Sharma et al. 2019). Recent analyses of FMRP binding sites in neurons found that Fmr1 preferentially associates with the coding regions of long mRNAs (Sawicka et al. 2019; Li et al. 2020).

Length has also emerged as an influential factor in the non-specific association of mRNAs with RNP particles. Repressive RNPs, including stress granules and heat shock granules, are significantly enriched for long mRNA transcripts (Padrón et al. 2019; Khong et al. 2017). The average length of stress granule-enriched transcripts is about ∼2-fold longer than non-enriched transcripts (Khong et al. 2017). Repressive particles are thought to form through multivalent interactions among and between proteins and RNA molecules (Molliex et al. 2015). Longer mRNAs contain more potential sites for promiscuous interactions with non-specific RNA binding proteins, including translational repressors, and with other RNA molecules. Therefore, an RNA granule-based system of gene regulation may inherently bias repressive effects towards genes encoding longer transcripts. It remains unclear if the two-fold boost FMRP imparts to the translation of large Drosophila mRNAs is related to their greater susceptibility to a similar level of non-specific repression within stress granules.

Here we have addressed whether FMRP truly acts differently in different species or tissues. By reanalyzing ribosome profiling data using a common pipeline from experiments conducted with FMRP-deficient Drosophila oocytes (Greenblatt and Spradling 2018) or mouse cortex (Das Sharma et al. 2019), we find that in both systems FMRP has an ancient, conserved function to promote rather than repress translation from a subpopulation of transcripts encoding large proteins. By analyzing the distribution of ribosomes along the length of FMRP target genes, we find that ribosome footprints of FMRP targets are not reduced due to an alleviation of ribosome stalling as would be expected from the FMRP repressor model.

Consistent with FMRP’s function as an activator, we find that many FMRP ASD target genes are haploinsufficient but not triplosensitive. We further show that ASD associated-genes encode mRNAs that are much longer than average, even when compared to other neuronally-expressed transcripts, explaining FMRP’s preferential association with ASD/ID targets.

Additionally, we analyzed the RNP particles that form between FMRP and one of its longest target mRNAs. About 35% of mRNAs encoding the 590 kDa E3 ubiquitin ligase Poe/UBR4 were detected within RNP particles containing both FMRP and Poe/UBR4 protein. Poe/UBR4 constitutes one of the 7 UBR box family E3 ligases, and together with UBR1, UBR2, and UBR5 mediate the N-end rule pathway of protein turnover (Varshavsky 1996; Tasaki et al. 2009; Varshavsky 2019). Our findings suggest that Poe/UBR4 particles form in response to the association of Poe/Ubr4 mRNA with FMRP and engaged ribosomes, and are sites of active Poe/UBR4 translation. Poe/UBR4 acts as a scaffold to recruit the E2 ubiquitin conjugating enzyme Rad6 and the RING E3 ligase KCMF1 to N-end rule substrates to activate their degradation (Hong et al. 2015a). This process is important in some Drosophila peripheral nerve paraneurial glial cells (Yager et al. 2001; Zülbahar et al. 2018) and myofibers (Hunt et al. 2019) to prevent excessive growth, suggesting a mechanism connecting the loss of FMRP and the downregulation of Poe/UBR4 to changes in growth signaling that could impact neural development.

## RESULTS

### FMRP has an ancient, conserved function to promote the translation of large proteins

In order to test whether FMRP has a similar function in Drosophila oocytes and in mouse cortex, we compared the effects of Fmr1 knockdown (KD) on translation as measured using ribosome profiling in these two tissues and organisms (Greenblatt and Spradling 2018; Das Sharma et al. 2019) but re-analyzed here (Table S1) from primary reads with identical bioinformatic pipelines (see Methods). Consistent with Greenblatt et al., we found that many transcripts containing long CDS were translationally downregulated in Fmr1 RNAi oocytes as compared to controls (Fig. 1A). The median CDS length of genes whose translation decreased significantly in Fmr1 KD oocytes was 3.1-fold longer than the median length of all oocyte expressed genes (4262 bps vs. 1372 bps respectively, Fig. 1B). We observed a strikingly similar effect of Fmr1 KD in the mouse cortex. Many genes encoding large proteins were translationally downregulated over a similar range (Fig. 1C), and the affected mRNAs had 3.4-fold longer CDS lengths as compared to all cortex-expressed genes (4797 bps vs 1401 bps respectively, Fig. 1D).

**Figure 1.**
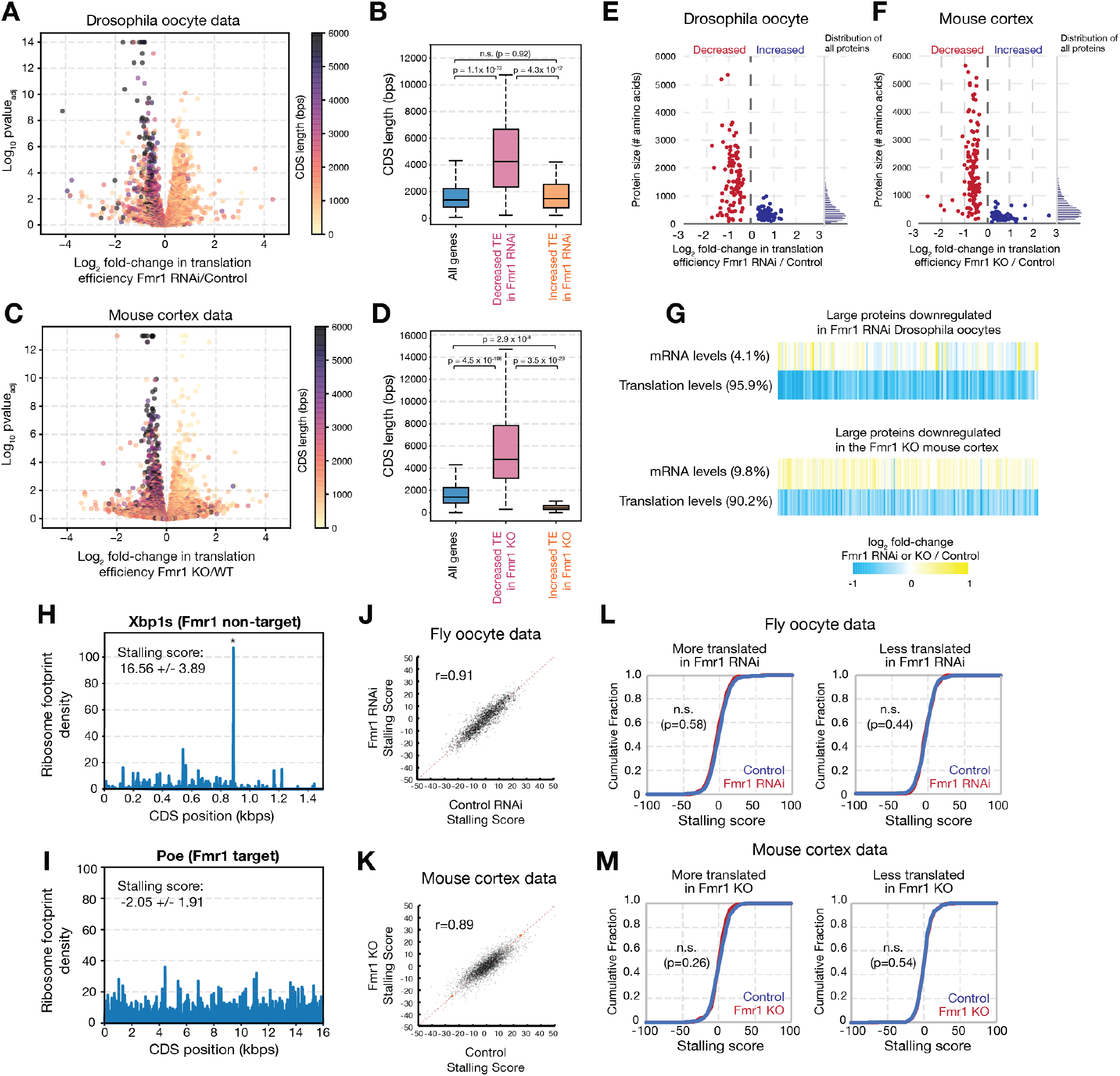
Reduced translation of large proteins in Fmr1 RNAi Drosophila oocytes and Fmr1 KO mouse cortex. (A–D) Volcano and box plots showing concordantly diminished translation efficiency of mRNAs with long CDS regions in Drosophila oocytes (Greenblatt and Spradling 2018) (A,B) and in mouse cortex (Das Sharma et al. 2019) (C,D). (E, F) Scatter plots showing reduced translation efficiency as a function of protein size for the 200 most significantly affected genes in Fmr1 RNAi Drosophila oocytes (E), or Fmr1 KO mouse cortex (F). Histograms showing size distributions for all Drosophila or mouse proteins respectively are shown to the right. (G) Heat map showing concordantly reduced translation levels but not mRNA levels for genes with reduced translation efficiency encoding large proteins >1,000 amino acids. Percentages reflect the fraction of genes in which translation efficiency changes were dominated by altered mRNA or translational levels. (H,I) Stalling scores (see Methods) were computed from the Drosophila oocyte ribosome profiles of the non-Fmr1 target Xbp1 (H) and the Fmr1 target Poe/Ubr4 (I). (J, K) Scatter plots showing high Pearson’s correlation of stalling scores from genes expressed in controls vs. Fmr1 RNAi oocytes (J) or the Fmr1 KO mouse cortex (K). Cumulative distribution plots showing no statistically significant changes in the stalling scores for genes that were translationally upregulated or downregulated in Fmr1 RNAi Drosophila oocytes (L) or the Fmr1 KO mouse cortex (M) (Kolmogorov-Smirnov test).

FMRP contains multiple RNA binding domains and studies suggest it has the potential to bind to widespread tetranucleotide and G-rich sequences (Anderson et al. 2016; Vasilyev et al. 2015; Ascano et al. 2012). Thus, FMRP-dependent translational activation might depend on length, with longer mRNAs being more strongly affected on average simply due to a greater number of FMRP binding sites. An alternative possibility is that FMRP translational activation requires only that a minimum number of binding sites be occupied by FMRP, reflecting a highly cooperative (and saturable) process. This latter model predicts that FMRP translational activation would be independent of mRNA length, at least beyond a threshold (i.e. the length at which the minimum number of binding sites are occupied). Our data were strongly consistent with the threshold model in both Drosophila oocytes or the mouse cortex. While genes translationally downregulated in Fmr1-deficienct cells were highly enriched for those encoding large proteins, the magnitude of the reduction in translation did not increase proportionately with CDS length (Fig. 1E,F). Changes in mRNA levels played little or no role in the reduced translational efficiency of large proteins in FMRP-deficient cells (Fig. 1G).

### FMRP does not affect translation by changing ribosome elongation or stalling rates

Declines in ribosome footprints on target mRNAs in FMRP-deficient tissues could in theory result from reduced translation initiation or increased elongation relative to controls. Drosophila oocytes and neurons are known to contain lower levels of three large FMRP target proteins in FMRP-deficient cells, consistent with reduced translation initiation (Greenblatt and Spradling 2018; Sears et al. 2019). However, ribosome density was reported to be more uniform across translated mRNAs in FMRP-deficient neurons (Das Sharma et al. 2019), a finding potentially consistent with a role for FMRP in translation elongation.

The presence of a ribosome pause site may potentially affect translational output only if translation elongation is rate-limiting. We surmised that if translation elongation is rate-limiting due to ribosomal stalling, then this should lead to a 5’ bias in the abundance ribosome footprints, with more reads aligning to the 5’ ends of coding sequences than the 3’ ends. The lack of read asymmetry in ribosome profiling data was used previously to argue that translation in vegetative yeast is normally controlled by initiation, with elongation rates having a negligible impact on protein production (Erdmann-Pham et al. 2020). In contrast, stalls impacting translation can be induced in bacteria lacking the factor EFP, which is required for the rapid translation of polyproline tracts (Doerfel et al. 2013; Ude et al. 2013), leading to asymmetry in ribosome footprinting profiles (Woolstenhulme et al. 2015).

We applied a “stalling score” metric (see Methods) to determine whether the presence of a stall site(s) led to decreased translation, as evidenced by asymmetry in the abundances of ribosome footprints aligned to either the 5’ or 3’ end of transcripts’ CDS regions. Large positive stalling scores would represent a build-up of reads at the 5’ end of the CDS region as compared to the 3’ end, consistent with a significant stalling effect leading to reduced translation; a negative stalling score would represent a build-up of reads at the 3’ end. For example, Xbp1, which has an internal ribosomal stall site essential for its membrane targeting, has a stalling score of 16.56 +/-3.89, reflecting the greater abundance of ribosome footprints prior to the stall as compared to after the stall (Fig. 1H). By contrast, the validated Drosophila Fmr1 target Poe/Ubr4 (Greenblatt and Spradling 2018) does not have a noticeable ribosomal stall peak, and consistent with this, Poe/Ubr4 has a low stalling score -2.05 +/-1.95 (Fig 1I).

In order to determine whether the changes in ribosome footprint abundance of Fmr1 target genes are significantly influenced by ribosome stalling, we compared stalling scores of genes in either control or Fmr1-deficient oocytes and neurons. Stalling scores were little affected by Fmr1 KO, since the scores strongly correlated genome-wide comparing wild type vs. Fmr1 RNAi Drosophila oocytes, as well as the mouse cortex from control vs. Fmr1 KO animals (Pearson’s r = 0.93 and r = 0.89 respectively, Fig. 1J,K). Inconsistent with the model that the reduced levels of ribosome footprints in Fmr1 targets are due to alleviated ribosome stalling, genes that were translationally downregulated or upregulated in the Drosophila oocytes and in the mouse cortex had statistically identical distributions of stalling scores (Fig. 1L,M). These data strongly argue that changes in ribosomal stall sites in Fmr1 RNAi or KO animals do not impact the overall translation of either Fmr1 targets or non-targets.

### Underproduction rather than overproduction of FMRP ASD target genes leads to clinical phenotypes

The relatively modest decreases in protein production that result from loss of Fmr1 function might be particularly significant in the case of “dosage sensitive” genes that cause defects when present in only one instead of two genomic copies. Dose sensitive genes relevant for the FXS phenotype, based on the FMRP activator model, would be “haploinsufficient” i.e. loss of either the maternal or paternal copy would lead to clinical defects. In contrast, under the repressor model relevant FMRP targets would be “triplosensitive,” where gene duplications leading to the overexpression of FMRP targets result in clinical phenotypes. We identified 61 genes that are translationally downregulated in the Fmr1 KO mouse cortex, and that score as SFARI Class I, Class II, or syndromic ASD genes (Table 1), which we considered to be potentially relevant. From this group, haploinsufficiency and triplosensitivity scores were curated in the Clinical Genome Resource (NIH) database from clinical data for 32 FMRP targets (Rehm et al. 2015). Consistent with the activator model, we found that 22 of the 32 downregulated ASD genes were classified as haploinsufficient at the levels of “emerging evidence” and “sufficient evidence” (score of 2 and 3 respectively, Table 1). In contrast, and in contradiction to the FMRP repressor model, none of these genes (0/32) had emerging or sufficient evidence of triplosensitivity. These data indicate that reduced expression, but not overexpression, of FMRP ASD targets lead to observable clinical phenotypes, and further support our model that FMRP acts primarily as an activator and not a repressor of gene expression.

**Table 1.** SFARI Class I and Class II ASD genes are translationally reduced in the Fmr1 KO mouse cortex. Table showing SFARI Class I and Class II ASD genes that are significantly altered in their translation efficiency in the Fmr1 KO mouse brain. 98% (61/62) of altered ASD genes are translationally reduced. ClinGen Dosage scores (3 = sufficient evidence, 2 = emerging evidence, 1 = little evidence, 0 = no evidence) (Rehm et al. 2015) show that many of these ASD genes have strong or emerging evidence of human syndromes associated with gene losses (haploinsufficiency) but not gene duplications (triplosensitivity).

| SFARI<br>Autism<br>Genes | Fold-change TE<br>Fmr1 KO / WT | p-valueadj | # Amino<br>acids | Darnell et al.<br>2011 CLIP<br>Target | ClinGen Haplo-<br>insufficiency Score | ClinGen Trip<br>sensitivity Sc |
| --- | --- | --- | --- | --- | --- | --- |
| PRR12 | 0.51 | 0.0E+00 | 2,035 | ✓ | n/a | n/a |
| KMT2A | 0.58 | 9.0E-05 | 3,966 | ✓ | 3 | 0 |
| HIVEP3 | 0.58 | 4.7E-04 | 2,348 | ✓ | 1 | 0 |
| SHANK1 | 0.59 | 0.0E+00 | 2,167 | ✓ | 1 | 0 |
| SETD1B | 0.59 | 1.4E-03 | 1,985 |  | n/a | n/a |
| KMT2C | 0.61 | 4.0E-08 | 4,904 | ✓ | 3 | 0 |
| SPEN | 0.63 | 2.7E-07 | 3,620 | ✓ | 1 | 0 |
| MED13L | 0.63 | 4.8E-08 | 2,207 | ✓ | 3 | 1 |
| TCF20 | 0.64 | 7.1E-07 | 1,987 | ✓ | 3 | 0 |
| SCAF4 | 0.64 | 6.1E-04 | 1,209 |  | n/a | n/a |
| AHDC1 | 0.65 | 2.2E-07 | 1,594 | ✓ | 3 | 0 |
| SRCAP | 0.65 | 6.2E-07 | 3,271 |  | 1 | 0 |
| CIC | 0.66 | 5.4E-07 | 1,604 | ✓ | 2 | 0 |
| KDM3B | 0.66 | 1.3E-06 | 1,762 |  | n/a | n/a |
| CNOT3 | 0.66 | 5.0E-03 | 751 |  | 0 | 0 |
| MAP1A | 0.67 | 0.0E+00 | 2,776 | ✓ | n/a | n/a |
| SHANK2 | 0.67 | 5.9E-08 | 1,472 | ✓ | 2 | 0 |
| RIMS1 | 0.67 | 9.7E-07 | 1,190 |  | 1 | 0 |
| AUTS2 | 0.67 | 8.0E-06 | 1,261 | ✓ | 3 | 0 |
| RERE | 0.68 | 1.0E-03 | 1,558 | ✓ | n/a | n/a |
| SYN1 | 0.68 | 1.2E-06 | 670 | ✓ | 2 | 0 |
| HECTD4 | 0.68 | 0.0E+00 | 4,418 |  | n/a | n/a |
| LRP1 | 0.69 | 0.0E+00 | 4,545 | ✓ | n/a | n/a |
| CAMK2A | 0.69 | 3.0E-04 | 478 | ✓ | 1 | 0 |
| CREBBP | 0.69 | 1.1E-06 | 2,441 | ✓ | 3 | 0 |
| NOVA2 | 0.70 | 2.6E-03 | 492 |  | n/a | n/a |
| CHD8 | 0.70 | 2.7E-05 | 2,582 | ✓ | 3 | 0 |
| SHANK3 | 0.71 | 5.4E-03 | 1,805 | ✓ | 3 | 0 |
| ARID2 | 0.71 | 7.3E-04 | 1,828 | ✓ | 3 | 0 |
| ASH1L | 0.71 | 2.9E-09 | 2,958 | ✓ | 3 | 0 |
| IQSEC2 | 0.71 | 1.3E-05 | 1,479 | ✓ | 3 | 0 |
| ZC3H4 | 0.72 | 3.6E-03 | 1,255 | ✓ | n/a | n/a |
| SON | 0.72 | 5.2E-04 | 2,444 | ✓ | 3 | 0 |
| GRIK5 | 0.72 | 5.6E-03 | 979 | ✓ | n/a | n/a |
| CACNA1A | 0.72 | 1.6E-05 | 2,327 | ✓ | 3 | 0 |
| KIF5C | 0.73 | 1.4E-10 | 956 | ✓ | n/a | n/a |
| HUWE1 | 0.73 | 6.6E-10 | 4,378 | ✓ | 0 | 0 |
| ZMIZ1 | 0.74 | 1.5E-03 | 1,066 | ✓ | n/a | n/a |
| AKAP9 | 0.74 | 1.2E-03 | 3,779 | ✓ | n/a | n/a |
| NCOR1 | 0.74 | 4.1E-07 | 2,454 | ✓ | n/a | n/a |
| ARID1B | 0.75 | 1.8E-04 | 2,244 | ✓ | 3 | 0 |
| AGO2 | 0.76 | 1.2E-03 | 860 |  | n/a | n/a |
| MYT1L | 0.77 | 6.6E-03 | 1,185 | ✓ | 3 | 0 |
| FOXG1 | 0.78 | 8.6E-03 | 481 |  | 3 | 0 |
| MECP2 | 0.78 | 9.5E-03 | 484 |  | n/a | n/a |
| WDFY3 | 0.78 | 6.5E-06 | 3,508 | ✓ | n/a | n/a |
| VAMP2 | 0.78 | 6.7E-05 | 116 | ✓ | n/a | n/a |
| NSD1 | 0.79 | 4.3E-03 | 2,691 | ✓ | 3 | 0 |
| CHD3 | 0.79 | 4.8E-04 | 2,055 | ✓ | n/a | n/a |
| EP400 | 0.80 | 8.2E-04 | 2,999 | ✓ | n/a | n/a |
| INTS1 | 0.80 | 5.2E-03 | 2,222 | ✓ | n/a | n/a |
| PHF3 | 0.80 | 2.7E-03 | 2,025 |  | 1 | 0 |
| YWHAG | 0.80 | 5.8E-04 | 247 | ✓ | n/a | n/a |
| SLC12A5 | 0.81 | 5.0E-03 | 1,138 | ✓ | n/a | n/a |
| CTNND2 | 0.81 | 2.3E-04 | 1,221 | ✓ | 2 | 0 |
| SATB1 | 0.82 | 7.4E-03 | 764 |  | n/a | n/a |
| TRRAP | 0.82 | 2.0E-04 | 3,847 | ✓ | 1 | 0 |
| NFIX | 0.82 | 5.5E-04 | 433 | ✓ | n/a | n/a |
| SYNE1 | 0.82 | 4.1E-03 | 1,431 | ✓ | n/a | n/a |
| TANC2 | 0.83 | 7.6E-03 | 1,994 | ✓ | n/a | n/a |
| ATP2B2 | 0.83 | 5.5E-04 | 1,198 | ✓ | n/a | n/a |
| ILF2 | 1.25 | 9.5E-03 | 390 |  | n/a | n/a |

FMRP target ASD genes were translationally downregulated to 51-83% of control levels based on reduced ribosome footprints. On average, FMRP target ASD genes were translationally reduced to similar levels as compared to generic downregulated genes (72% vs. 70% respectively). FMRP probably acts by directly binding to most of these mRNAs, since 48 of 61 (79%) were cross-linked in FMRP CLIP experiments (Darnell et al. 2011). Thus, at least 22 dosage sensitive ASD genes are translationally downregulated in the Fmr1 cortex, likely through the loss of FMRP direct binding, each potentially contributing to defects despite relatively small individual reductions.

### ASD/ID genes encode exceptionally large proteins whose requirement for Fmr1 is evolutionarily conserved

FMRP is known to associate with mRNAs from many genes associated with ASD and ID (Darnell et al. 2011; Sawicka et al. 2019). To investigate the average size of ASD/ID genes and their encoded proteins, we compared them with the general population of neuronally-expressed genes in the juvenile mouse cortex (Das Sharma et al. 2019). We classified SFARI Class I and Class II autism genes (Abrahams et al. 2013; Pereanu et al. 2017) as “ASD/ID genes,” and found they encode proteins averaging 1,234 amino acids, compared to 589 amino acids on average for neuronally-expressed genes. There was an even greater average difference in the size of these genes, 83kb vs 23 kb, consistent with prior studies (King et al. 2013; Zhao et al. 2018). The greater size was largely because ASD/ID genes contain larger introns totaling 78.3 kb on average vs. 20.6 kb on average for neuronally expressed genes (Fig. 2B). ASD/ID genes were only slightly larger than the neuronal average with respect to 5’ UTR length (250 bp vs. 160 bp, Fig. 1C), and 3’ UTR length (1703 vs. 1053 bp, Fig. 2D). ASD/ID genes were not significantly different from neuronal genes in mRNA levels (Fig. 2E), or translation levels (Fig. 2F).

**Figure 2.**
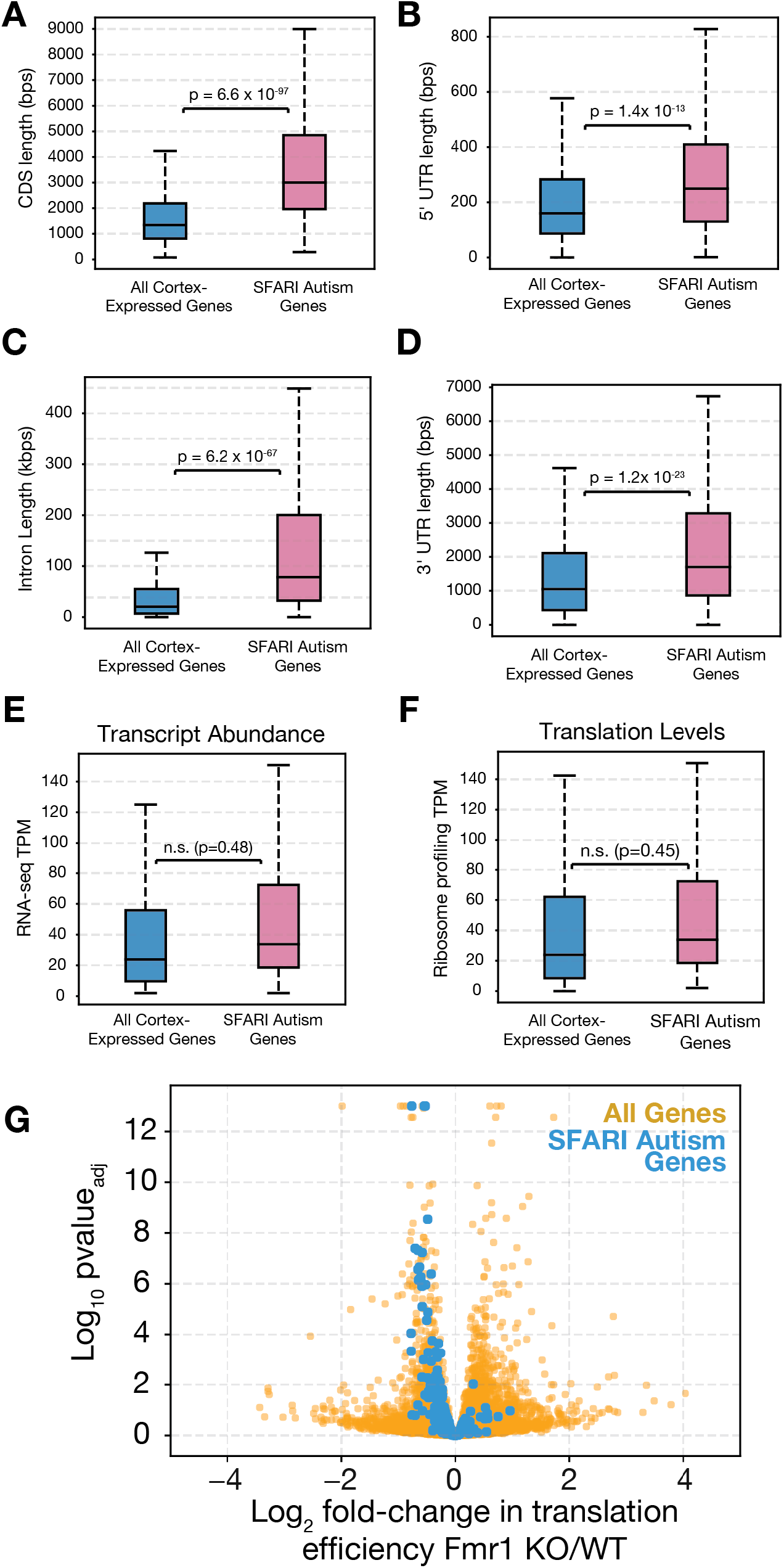
SFARI autism genes have longer open reading frames, UTRs and introns than average. Box plots of length distributions showing increased (A) coding sequences lengths, (B) 5’ UTR lengths, (C) intron lengths (summed across all introns), and (D) 3’ UTR lengths specifically for SFARI class I and class II autism spectrum disorder (ASD) genes (magenta) as compared to all genes expressed in the juvenile mouse cortex (blue). (E, F) No significant differences were detected between the distributions of transcript levels (E) or translation levels (F) specifically for ASD genes vs. all cortex-expressed genes as detected by mRNA sequencing and ribosome footprinting respectively. (G) Volcano plot showing a concordant downregulation in the translation efficiency of SFARI Class I and Class II autism genes (blue) as compared to all neuronally expressed genes (orange).

Not surprisingly given their large average mRNA length, ASD/ID genes as a class were translationally downregulated in the FMRP KO cortex (Figure 2G). FMRP target ASD genes encode proteins even larger than the average for ASD genes generally (2,035 aa vs 1,234 aa). However, the presence of ASD/ID genes as FMRP targets is probably not due to size alone, since a significant fraction of ASD/ID genes controlled by FMRP in mouse cortex and Drosophila oocytes (22/61, Table S2) are orthologs (p-value = 2.0 × 10^−28^, chi-squared test).

Thus, FMRP-dependent ASD/ID genes most likely carry out functions in nervous tissue and oocytes that require much larger and more dose-sensitive proteins than average.

### Poe particles are likely sites of FMRP-dependent translational activation

FMRP is associated in neurons and oocytes with some RNP granules that contain markers of P bodies and stress granules (Rosario et al. 2016; Barbee et al. 2006). These structures normally contain mRNAs that are relatively inactive in translation. In contrast, electron microscopic analyses reveal a close association of FMRP-containing particles with ribosomes (Antar et al. 2005). Consequently, we sought to learn more about the nature, diversity and function of RNP particles containing FMRP in Drosophila follicles, and in particular to address whether they are sites of active translation rather than repression.

We focused on a previously identified granule (“Poe particle”) containing FMRP target protein Poe/UBR4 located in the nurse cells of Drosophila ovarian follicles (Greenblatt and Spradling 2018). We co-stained nearly mature (stage 10) follicles with antibodies to Poe and to FMRP, and found that Poe/UBR4 particles are highly enriched for FMRP (Fig. 3A). To test whether Poe mRNA is translated in Poe particles, we investigated whether full-length Poe/UBR4 polypeptides (and not just nascent fragments) are present in Poe particles. We generated C-terminally tagged Poe/UBR4-GFP using CRISPR-mediated homologous recombination. We found that GFP-tagged full-length Poe/UBR4 (“Poe-GFP”) proteins are enriched in Poe particles (Fig. 3B). Using single molecule FISH (smFISH) labeling of Poe/UBR4 mRNA molecules, we found that Poe/UBR4 mRNA, but not the FMRP non-target Dhc64C mRNA, is also enriched in Poe particles (Fig. 3C). We documented co-localization using line scans (Fig. 3C) as described in Methods. Thus, multiple Poe/UBR4 mRNAs are recruited to FMRP-containing Poe particles, consistent with the idea that high translation levels in these particles contribute to FMRP-dependent Poe translational activation. We hypothesized that Poe particles, and possibly other FMRP-containing particles, are sites where translation of FMRP targets is activated, leading to higher translation output than in the absence of FMRP. In this view, the lower levels of Poe in Fmr1 KD oocytes would be due to the loss of Poe mRNA translation in FMRP-containing Poe particles, and their derivatives.

**Figure 3.**
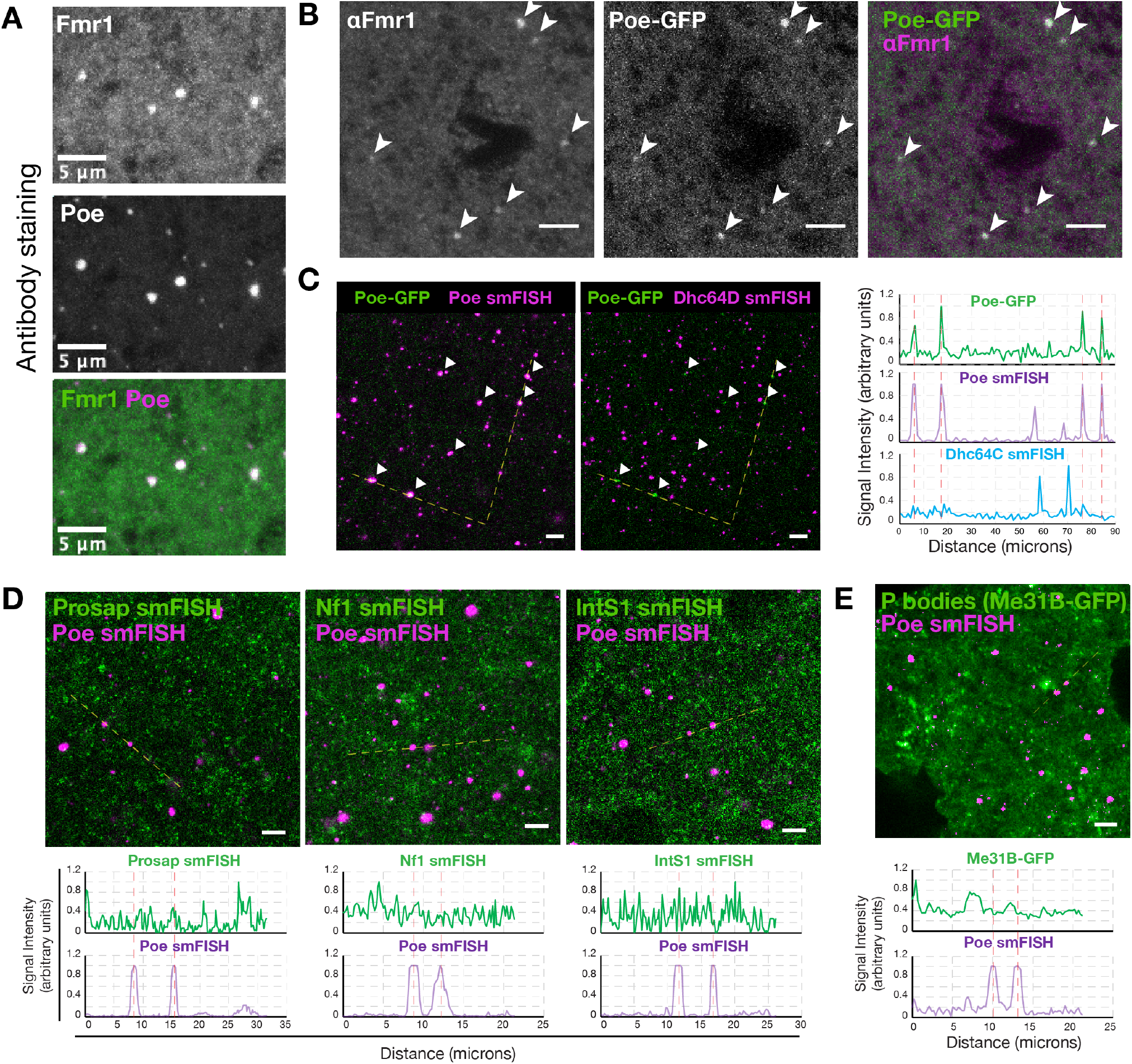
FMRP–Poe/UBR4 particles are putative sites of Poe/UBR4 translation. (A) Immunostaining shows co-localization of FMRP and Poe protein in particles in the nurse cells of stage 10 egg chambers. (B) Immunostaining showing that full-length Poe protein (Poe-GFP) is enriched in Poe-FMRP particles. (C) smFISH showing that Poe mRNA, but not Dhc64C mRNA, is enriched in Poe protein particles. (right) Signal intensities (along drawn dotted yellow dashed line) of Poe-GFP and Poe smFISH, but not Dhc64C smFISH, are highly correlated. (D) smFISH and line scans (along drawn dotted yellow dashed lines) showing that other FMRP target mRNAs, Prosap, Nf1, and IntS1 are not enriched in Poe particles. (E) smFISH and line scan showing that Poe particles are independent of the P body marker Me31B-GFP.

We next investigated whether other FMRP target mRNAs are enriched in FMRP-Poe particles. We labeled three additional FMRP target mRNAs, Nf1, Prosap, and IntS1, using smFISH (Fig. 3D). These mRNAs are distributed widely in the cytoplasm and form particles smaller than the larger Poe particles but do not co-localize with Poe particles (Fig. 3D). Poe particles are distinct from P bodies, structures which are thought to be enriched for translationally repressed mRNAs, since they do not co-localize with the P body marker ME31B (Fig. 3E). Together, these data indicate that at least one target of FMRP-enhanced translational initiation, Poe, is associated with FMRP in a distinct particle, but FMRP-Poe particles are not general sites for FMRP target translation.

### The presence of FMRP-Poe RNP particles marks FMRP-dependent translational activation

Poe particles form in stage 10 follicles as growth slows and follicles transition toward maturation and quiescence (Greenblatt and Spradling, 2018). In stored oocytes FMRP activates the translation of Poe/Ubr4 by ∼2-fold without affecting Poe/Ubr4 mRNA levels (Greenblatt and Spradling, 2018). In contrast, FMRP had no effect on the translation of another very large protein of similar size (∼530 kDa), Dhc64C (Fig. 4A). FMRP was required for the formation of Poe particles containing Poe mRNA, because unlike wild type follicles, Poe particles were absent and Poe mRNA was dispersed in follicles lacking FMRP (Fig. 4B,B’). Dispersed Poe mRNA could still be detected above background, because cytoplasmic Poe smFISH signal was completely absent from egg chambers expressing Poe RNAi (Fig. 4B,B’).

**Figure 4.**
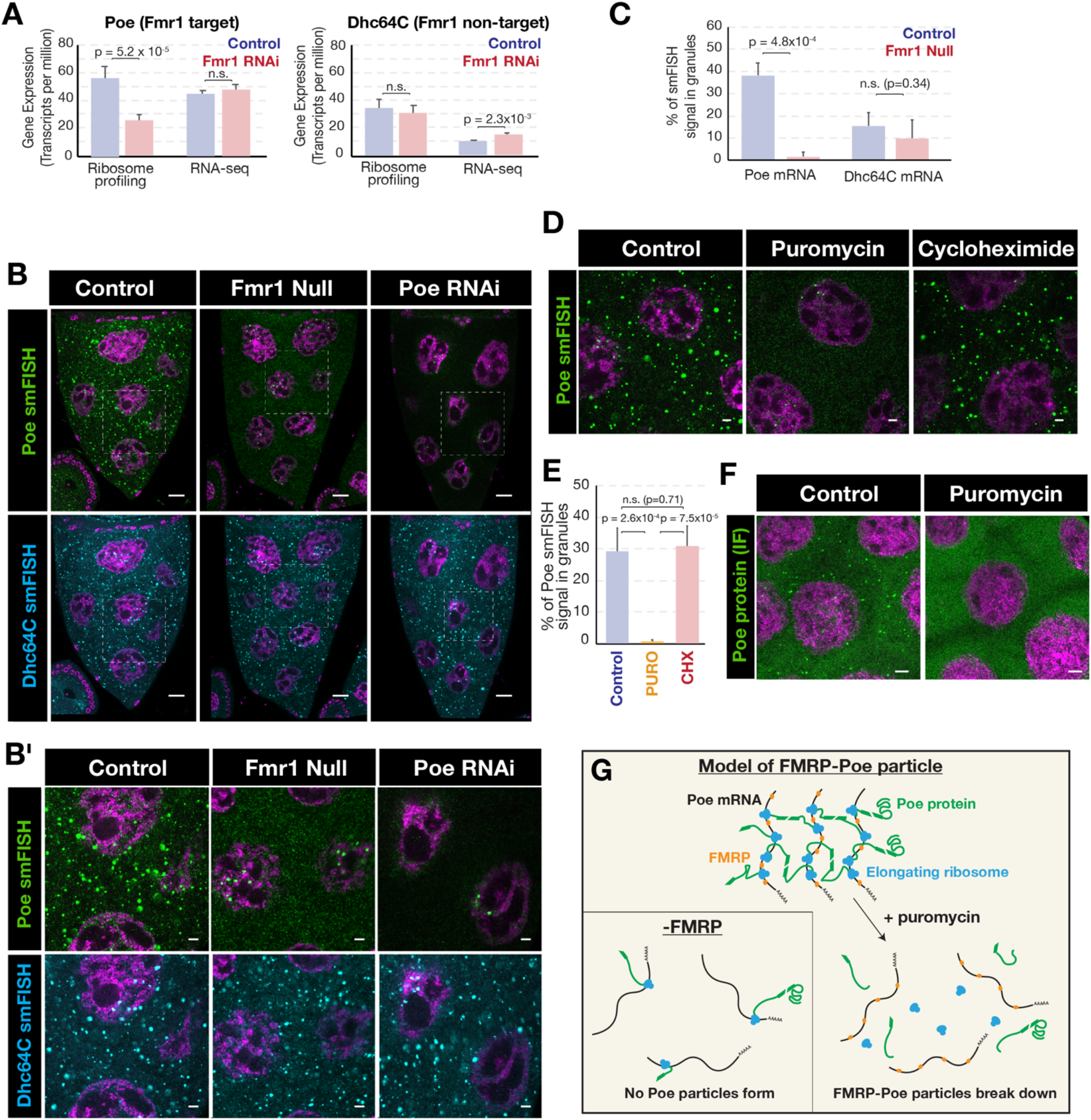
The presence of FMRP-Poe/UBR4 particles is a sensitive readout of FMRP-dependent translation. (A) Plots showing that the translation of Poe mRNA, but not another large mRNA Dhc64C, is reduced in Fmr1 RNAi oocytes. (B, B’) smFISH showing that Poe mRNA and Dhc64C mRNA are localized to large particles in wild type egg chambers. Fmr1 RNAi results in the dispersal of cytoplasmic Poe mRNA signal from particles, but not Dhc64C mRNA or nuclear Poe mRNA signal (presumed transcription sites). The dispersed cytoplasmic Poe mRNA smFISH signal is specific as it is eliminated in Poe RNAi egg chambers. (C) Plots of image analyses (see Methods), showing that recruitment of Poe mRNAs to particles is largely eliminated in Fmr1 RNAi egg chambers, whereas there is no significant effect on recruit of Dhc64C mRNA to particles. (D) smFISH showing that Poe mRNAs are rapidly dispersed upon premature translation termination with 15 mins. puromycin treatment, but with a similar treatment with the elongation inhibitor cycloheximide. (E) Quantification showing reduced localization of Poe mRNA to particles in puromycin-treated, but not cycloheximide-treated egg chambers. (F) Immunostaining of control or puromycin-treated egg chambers shows that Poe protein particles are also lost upon polysome assembly. (G) Model of FMRP-dependent translation of Poe in particles. High local concentrations of interaction domains contained with polysome-associated Poe protein, driven by FMRP translation activation, overcomes the entropy barrier to drive the formation of microscopically visible particles. Diminished translation in the absence of FMRP leads to the entropic dispersal of Poe mRNA and protein.

Dhc64C mRNA similarly formed microscopically visible particles in late follicles; however, unlike Poe particles, Dhc64C particles were unaffected by the loss of FMRP. From image analyses of smFISH-labeled egg chambers (see Methods) we estimated that 35% of Poe mRNA is normally found in particles in wild type egg chambers, but <2% in FMRP-null egg chambers (Fig. 4C). In contrast, ∼15**%** Dhc64C mRNA is present in DHC particles in wild type egg chambers, and this value was not changed in the absence of FMRP (Fig. 4C). These experiments show that FMRP is required for the formation of Poe particles but is not required generally for cytoplasmic RNA granule formation. Thus, the presence of Poe particles can serve as an indicator of ongoing FMRP-dependent translational enhancement activity.

### FMRP-Poe RNP particles contain actively translocating ribosomes, and disperse upon polysome disassembly

RNP granules are thought to form as a result of multiple weak interactions between RNA and RNA binding proteins (Matheny et al. 2020). In addition, protein interaction domains found in partially translated polypeptides can also direct polysomes to specific subcellular sites (Yanagitani et al. 2011). We hypothesized that if Poe particles are formed through weak interactions between FMRP, Poe mRNA, and partially translated Poe protein, then drug treatments which separate elongating protein chains from their mRNAs should result in the rapid loss of Poe particles. We treated wild type follicles briefly (15 minutes) with either puromycin, an analog of the 3’ end of tyrosol-tRNA, whose incorporation by actively elongating ribosomes leads to premature chain termination and polysome disassembly, or cycloheximide, which interferes with ribosomal translocation, thus blocking elongation and “freezing” polysome-associated ribosomes in place. A brief treatment of follicles with puromycin led to the rapid dispersal of Poe mRNA, phenocopying the effect of Fmr1 loss (Fig. 4D,E). In contrast, a similar treatment with cycloheximide did not change Poe mRNA distribution (Fig. 4D,E).

The puromycin sensitivity of FMRP-Poe particles also contrasts with prior *in vitro* data that FMRP-associated polysome complexes are translationally stalled and therefore should be puromycin resistant (Darnell et al. 2011). Puromycin treatment also resulted in a similar rapid dispersal of Poe protein (Fig. 4F), indicating that Poe protein remains in particles following local translation through its mRNA association.

These experiments support a simple model of Poe particles structure and formation (Fig. 4G). Poe particles form when nascent Poe chains are produced on FMRP-bound Poe mRNAs and self-associate across mRNAs. Some full-length Poe molecules also remain associated where they are able to mature into active E3 ligase complexes prior to release into the cytoplasm. The rapid dispersal of Poe protein and mRNA from particles following puromycin treatment suggests that Poe particles represent steady state structures with a high degree of flux, in which ongoing translation balances the release of completed Poe E3 ligases. Since dispersed but significant Poe translation remains in the absence of FMRP, however, the disappearance of Poe particles under these conditions suggests that net particle growth depends critically on a threshold density of Poe nascent chains which is only reached through FMRP activation (Fig. 4G).

## DISCUSSION

### Fmr1 has an ancient, conserved function to promote the translation of large proteins

Our side by side comparison of ribosome profiling data show that the effect of Fmr1loss on translation in Drosophila oocytes and mouse cortex is extremely similar despite the fact that different tissues and organisms are involved. Fmr1 activates rather than represses target mRNA translation in both systems. In both cases the targets are more than three times longer than average-sized mRNAs and encode correspondingly larger proteins, including many orthologs. In both cases, the normal stimulatory effects of FMRP are about two-fold or less, independent of the length of the mRNA and their predicted FMRP binding site content. Moreover, these similar effects are not due to changes in rates of ribosome elongation or stalling, but represent differences in translation initiation and protein output. Consistent with these findings, Western blotting documented the expected reductions in two FMRP target proteins in oocytes, the 2,053 amino acid snRNA 3’ processing factor IntS1 and the 5,322 amino acid E3 ubiquitin ligase Poe/UBR4, while the 3,719 amino acid protein kinase A regulator rugose/NBEA is reduced in *Fmr1* mutant Drosophila mushroom body (Sears et al. 2019).

These findings are consistent with the important and ancient role *Fmr1* plays in neurons, oocytes, and spermatocytes from diverse animals ranging from Drosophila to humans (Hagerman et al. 2017; Drozd et al. 2018). Mammalian and Drosophila FMRP contain the same protein domains and the expression of human FMRP rescues neural defects in Drosophila *Fmr1* mutants (Coffee et al. 2010). FMRP had long been considered primarily a translational repressor (Darnell et al. 2011; Laggerbauer et al. 2001; Li et al. 2001); however bulk translation in the brain may not be increased in the absence of *Fmr1* (Schmidt et al. 2020). Our findings here are consistent with observations that FMRP CLIP targets have concordantly reduced ribosome footprints (Das Sharma et al. 2019), and that FMRP preferentially binds to long transcripts (Sawicka et al. 2019; Li et al. 2020). Indeed, none of the mRNAs with increased translation in *Fmr1*-null animals were among the top 200 CLIP targets (Das Sharma et al. 2019; Darnell et al. 2011). The increased translation of some genes in Fmr1 mutant animals are likely due to secondary, indirect effects of Fmr1 loss. FMRP targets include transcription repressors, and E3 ubiquitin ligases that stimulate protein degradation via the proteasome, while some proteins increase in translation due to the dysregulation of mTOR signaling (Das Sharma et al. 2019; Sharma et al. 2010). These data help explain the failure of the mTOR inhibitor rapamycin to reverse behavior deficits in Fmr1 null mice (Saré et al. 2017), since treating the increased translation levels of secondary FMRP targets would not rescue the primary translational defect in these animals -the underproduction of dozens of ASD/ID-associated proteins. These data are also consistent with observations that underproduction, but not overproduction, of FMRP ASD targets leads to clinical phenotypes (Table 1).

### Fmr1 is used dynamically in neurons and other cells that depend on stored mRNAs

Large cells such as neurons and oocytes rely heavily on the transport and storage of stable, untranslated mRNAs. For example a large fraction of oocyte mRNAs are translationally repressed for later use during early embryonic development, including CNS development (Greenblatt et al. 2019; Kronja et al. 2014). Neurons transport translationally regulated synaptic mRNAs up to a meter or more down long axonal projections. Such transport requires cells to partition mRNAs into repressed or actively translated states through the use of RNP particles, including P bodies and neuronal particles, that are structurally related to stress granules.

These complexes often associate binding partners together through phase separation, which is sensitive to protein concentration (Molliex et al. 2015). Thus, there are periods when mRNAs undergoing transport from the cell body along an axon, or while stored in the vicinity of a synapse, must remain functional while in an inactive state. The rate of growth of Drosophila oocytes varies strongly in response to available nutrition, and both oocytes and neurons grow at very different rates at different stages of development. P bodies and stress granules can be seen to form and disperse within minutes when growth conditions vary sharply (Shimada et al. 2011; Wheeler et al. 2016). *Fmr1* function is particularly required under conditions where development is slowed, for example after mature oocytes enter quiescence (Greenblatt and Spradling 2018).

Previous observations showed that genes expressed during neural development on average encode larger proteins than in many other tissues (Gabel et al. 2015). Proteins regulated by *Fmr1* are more than three times larger still, and include many genes associated with autism and intellectual disability syndromes. There is currently no well understood functional explanation for what exceptionally large protein size contributes to higher neural functions. Neural development likely involves complex tasks such as integrating sensory inputs and outputs in ways that are vital to survival. It may be advantageous to place multiple protein domains along a single polypeptide chain in order to ensure they are present at equimolar levels to help guarantee the proper stoichiometries of protein domains that function together in a complex process. Another possibility is that large neural proteins function as scaffolds for exceptionally large and specific protein complexes that play critical functions essential for neural processing. Such proteins might require many critical binding sites for additional binding partners spaced substantial distances apart along the primary sequence. If processes associated with the most complex neuronal-based decision making are especially dependent on complexes catalyzed by enormous proteins, this might explain the enrichment of the ASD/ID gene products among the largest cohorts of FMRP targets.

### Identification of a specific RNP containing Ubr4/Poe, FMRP and ribosomes

What is the role of RNP particles in FMRP-dependent translational activation? It has become clear that under conditions of stress or growth quiescence, mRNAs frequently become packaged in repressive RNPs, such as stress granules or P bodies, potentially for reuse when conditions favoring activity return. Inactive particles may selectively incorporate large mRNAs for stochastic reasons, hence FMRP might have evolved to counteract this small but potentially deleterious force that would otherwise reduce the ongoing production of important large proteins. There are several ways FMRP might act to activate mRNA translation. FMRP binding at multiple sites along an mRNA might directly or with partner proteins counteract the equilibrium favoring its inclusion within repressive RNPs. Alternatively, bound FMRP might interact with linker and motor proteins to move target mRNAs to cytoplasmic domains more favorable for continued activity. Finally, FMRP binding along an mRNA might catalyze it to form novel particles that would stimulate increased translation. There may be FMRP target mRNAs that utilize one or more of all these mechanisms. Consequently, it was significant that we were able to characterize a specific oocyte RNP complex containing mRNA encoding the FMRP target Poe/UBR4, a highly conserved E3 ubiquitin ligase. Our study of Poe/UBR4 particles provided insight into how FMRP functions to enhance production of a particular target, Poe/UBR4, that may be of particular phenotypic significance.

The UBR family of E3 ligases have been shown to mediate N-end rule protein degradation (Varshavsky 2019). Seven families, Ubr1-Ubr7 are recognized, and all share a common zinc finger domain. UBR E3 ligases are thought to recognize substrates by their degrons, which are frequently generated by the activity of N-terminal asparagine or glutamine amido hydrolyases such as Ntan1 and Ntaq1 (tungas in Drosophila) and Ate1, although some degrons are present internally (Hwang et al. 2009). Unlike other UBR family members, UBR4 lacks a known ubiquitination domain (Tasaki et al. 2005) and rather acts as a scaffold to recruit E3 ubiquitin ligases to bound targets (Hong et al. 2015a). Together, this complex is able to recognize and process substrate proteins for degradation by the proteosome.

### Disruption of Poe/UBR4 particles may impact N-end rule proteolysis and cause neural defects

Poe particle formation may resemble the assembly of nuclear pores after mitosis, and in early Drosophila embryos. The translation of the nucleoporin protein Nup358 mRNA drives the formation of RNP particles enriched in NUP358 and other nucleoporins that serve as an “assembly platform” for the maturation of nuclear pore complexes (Hampoelz et al. 2019). FMRP-dependent translation of Poe mRNA may likewise generate assembly platforms for additional proteins involved in the N-end rule degradation pathway that subsequently attract and degrade their targets.

Knockdown of *Fmr1* or *Poe* in developing Drosophila female germ cells leads to oocytes with reduced capacity to support nervous system development (Greenblatt and Spradling 2018). Poe particles may be a major source of fully assembled complexes capable of catalyzing N-end rule mediated turnover. Studies of mutations in *Poe* and in upstream enzymes in the N-end rule pathway such as *Ntan1*, show that the regulation of MAPK and Hippo signaling is altered, possibly because *Poe*-mediated N-end rule protein degradation affects the levels of MAPK and Yorkie effectors (Zülbahar et al. 2018; Ashton-Beaucage et al. 2016). Loss of *Poe* is known to thicken the inner glial layer (“perineurium”) of Drosophila peripherial nerves through altered MAPK signaling, an effect further enhanced by disruption of Neurofibromatosis 1 (NF1) (Yager et al. 2001). Loss of the ability to generate a sufficient number of active Poe particles at appropriate times during development may be responsible.

### Poe particles may help detect FMRP-dependent translation and perturbations ameliorating target protein under-production

The formation of Poe/UBR4 particles offers a sensitive assay for identifying tissues undergoing Fmr1-mediated translational upregulation. Overall, the effects of Fmr1 are weak – a small fraction of mRNAs (∼5%) are FMRP targets and the increase in translation driven by FMRP is ∼2-fold or less. In contrast, the formation of FMRP–Poe particles provides a robust and sensitive readout of FMRP function; Poe/UBR4 particles are lost in Fmr1 mutants and recover if an extra wild type *poe* gene copies are provided (Greenblatt and Spradling 2018). By screening for factors that eliminate or increase Poe particles formation, it may be possible to identify factors important for the production of large proteins, as well as suppressors potentially capable of reversing the reduced translation phenotype in *Fmr1*-mutant cells.

It will be important to identify other FMRP target mRNAs that may work together. Two other N-end rule E3 ligase, UBR3 and UBR5 (known as *hyperplastic discs (hyd)* in Drosophila), also encode very large proteins, and appear to be FMRP targets (Table S1). UBR5 was identified as an FMRP CLIP target (Darnell et al. 2011), and is a SFARI group 3 autism gene candidate. Multiple additional E3 ligases are also FMR1 targets and associated with ASD/ID, including HUWE1, TRIP12, BIRC6, HECW2. The association of many transcripts encoded by ASD/ID genes with FMRP suggests that pharmacological interventions targeting this system could selectively promote the expression of dozens of genes individually associated with ASD/ID, and therefore may represent one means in which to at least partially rescue the expression of genes leading to syndromes caused by haploinsufficiency.

## METHODS

### Read alignments and quantification

Raw sequencing data from (Greenblatt and Spradling 2018) and (Das Sharma et al. 2019) were aligned to the Flybase Consortium/Berkeley Drosophila Genome Project (BDGP)/Celera Genomics Drosophila release dm 6.28 and Genome Reference Consortium mouse build 38 mm10 assemblies respectively. Alignments were performed with STAR (Dobin et al. 2013) and TPM values computed using RSEM (Li and Dewey 2011) for mRNA sequencing experiments. TPM values for ribosome profiling data were computed using featureCounts from the Subread (Liao et al. 2014) package for ribosome footprints aligning to CDS regions.

### Length and expression analyses

SFARI Class I and Class II genes were obtained from the SFARI Gene database (Abrahams et al. 2013). For each gene, the highest expressed transcript isoform was used as the representative form. Transcripts with a TPM value of <2 were excluded from analyses. Intron, UTR, and CDS lengths for transcript isoforms were computed from BDGP and refGene gene models for Drosophila and mouse data respectively. For translation efficiency analyses, fold-change and p_adj_ values were computed using RiboDiff software (Zhong et al. 2017). Stress-granule enrichment data were obtained from (Khong et al. 2017).

### Ribosome stalling analysis

Ribosome footprints were mapped to their P sites with a 3’ offset of 14 bases using the make wiggle function from the Plastid software package (Dunn and Weissman 2016). Ribosome footprinting read densities mapping to the 5’ and 3’ ends (first and last quintile) of transcript CDS regions were quantified with Plastid. The stalling scores for representative transcripts were computed as the (5’ CDS read density – 3’ CDS read density) / (sum CDS read density) x 100, where 5’ CDS and 3’ CDS read densities represent the sum of reads coming from the 5’-most and 3’-most 20% of the CDS region respectively. Significantly upregulated and downregulated genes were defined as genes whose TE increased or decreased in Fmr1 mutant or RNAi / control with a p_adj_ value of < .01. For genome-wide stalling analysis, standard deviations of stalling scores from either control of Fmr1-deficient replicates were computed. The quality of the stalling scores were determined by taking the minimum stalling score to standard deviation ratio for control and Fmr1 KO/RNAi experiments for each transcript. Transcripts with the top 4,000 stalling quality scores (i.e. transcript stalling scores with the least noise) were used for genome-wide analyses.

### Immunostaining and single molecule fluorescence *in situ* hybridization (smFISH)

Adult female flies were fed for 2-4 days on wet yeast paste and ovaries hand dissected in Grace’s Insect Medium (Life Technologies). Ovaries were fixed in 4% formaldehyde (37% formaldehyde diluted in PBST (0.2% BSA, 0.1% Triton X-100 in 1X PBS)) for 12 or 15 minutes for immunostaining and smFISH experiments respectively. For immunostaining, ovaries were incubated with primary antibodies diluted in PBST with gentle agitation overnight at 4°C . Ovaries were then washed 3 times in PBST for at least 20 min and incubated with secondary antibodies overnight. Ovaries were then washed 3 times with PBST for at least 20 min each, and DAPI (1:20,000-fold dilution of a 5mg/mL stock) was added to the last wash.

For smFISH experiments, fixed ovaries were washed three times in PBST for 10 min each, followed by one wash in Buffer A (Biosearch Technologies) for 20 min at room temperature. Samples were then incubated in freshly prepared hybridization buffer (Biosearch Technologies) for 30 min-2 hours at 37C. Samples were then incubated overnight at 37C in hybridization buffer containing smFISH probes (Biosearch Technologies) diluted 1:50. Samples were incubated in prewarmed Buffer A at 37C for 30 min, followed by three additional washes with Buffer A at room temperature. DAPI (1:20,000 dilution from a 5 mg/mL stock) was added to the final wash. Samples were then washed with Buffer B (Biosearch Technologies) for 10 minutes at room temperature.

### smFISH image acquisition and analysis

12 bit images were acquired with a Leica SP8 microscope using counting mode and 16 line accumulations. To calculate the fraction of mRNAs in particles, Gaussian blurred nuclei were computationally filtered out, and histograms of cytoplasmic smFISH signal were computed using ImageJ software (NIH). Images of particles were considered to be composed of “hot” pixels, identified as those containing >30 photon counts. The fraction of signal from particles was calculated as the fraction of signal coming from hot pixels as compared to the total cytoplasmic smFISH signal. Line scan intensity profiles were generated in ImageJ software.

## SUPPLEMENTAL MATERIAL

## TABLE LEGENDS

**Supplementary Table 1. Compiled RiboDiff output for Fmr1 RNAi Drosophila oocyte and Fmr1 KO mouse cortex analyses**.

**Supplementary Table 2. ClinGen haploinsufficiency scores for Drosophila orthologs of SFARI Class I, Class II, and syndromic autism genes translationally downregulated in oocytes**.

These tables are provided separately as Excel files.

